# Ethoscopes: an open platform for high-throughput *ethomics*

**DOI:** 10.1101/113647

**Authors:** Quentin Geissmann, Luis Garcia Rodriguez, Esteban J. Beckwith, Alice S. French, Arian R. Jamasb, Giorgio F. Gilestro

## Abstract

We present ethoscopes, machines for high-throughput analysis of behaviour in *Drosophila* and other animals. Ethoscopes provide a software and hardware solution that is reproducible and easily scalable; they perform, in real-time, tracking and profiling of behaviour using a supervised machine learning algorithm; they can deliver behaviourally-triggered stimuli to flies in a feedback-loop mode; and they are highly customisable and open source. Ethoscopes can be built easily using 3D printing technology and rely on Raspberry Pi microcomputers and Arduino boards to provide affordable and flexible hardware. All software and construction specifications are available at http://lab.gilest.ro/ethoscope.

## Introduction

Understanding how behaviour is coordinated by the brain is one of the ultimate goals of neuroscience. In particular, much of modern neurobiology focuses on finding the genes and the neuronal circuits underlying simple and complex behaviours alike, aiming to describe and eventually understand how the brain processes sensory inputs into motor outputs. For many years, starting from Seymour Benzer’s seminal work[1], the fruit fly *Drosophila melanogaster* has been considered one of the model organisms of choice to dissect the genetics of behaviour. In the past decade, *Drosophila* has also emerged as an excellent model for studying not only the genes, but the neuronal circuitry of behaviour too: the combination of a rapidly delineating connectome together with an unrivalled repertoire of genetic tools has established *D. melanogaster* as one of the most promising animal models to study neuronal circuits. Optogenetics, thermogenetics, a genome-wide collection of RNAi lines, and a plethora of crafted and carefully described GAL4 lines, constitute a robust arsenal for neurobiologists interested in studying the neuronal circuitry underpinning behaviour. The limiting factor for *ethomics* - the high-throughput approach to behavioural studies - is therefore not the availability of genetic tools, but rather the access to an objective, reproducible and scalable system to detect and classify behaviour. Historically, *Drosophila* neuroscientists have often shown a high degree of ingenuity in devising paradigms and creating apparati able to capture relatively simple behaviours in a high-throughput fashion, usually driven by the desire to perform genetic screens. Analysis of phototaxis[2], geotaxis[3], response to ethanol inebriation[4,5], olfactory learning and habituation[6,7], and biology of circadian behaviours[8] are all successful examples of clever paradigms that have allowed high-throughput screenings of specific behaviours. More recently, ad hoc solutions featuring computational approaches have also been introduced: some specifically dedicated to a subset of behaviours, such as sleep[9–11] or feeding[12,13], and others designed to be more versatile[14–17]. While computer-assisted analysis of behaviour has the potential to revolutionise the field, adoption and throughput of currently available techniques are limited by several factors. Above all, the requirement for a non-standardised hardware-setup, which often bears problems of cost, footprint and scalability. Typically, most other systems consist of a centralised system in which one or several cameras record high-resolution videos that are then processed, in real-time[9–11] or offline[14,16,18,19], by a central, powerful workstation. To lower entrance barriers to machine analysis of behaviour, we developed the ethoscope platform. In devising its architecture - decentralised and modular - we took inspiration from the commercially available Drosophila Activity Monitors (DAMs, TriKinetics Inc., Waltham, Massachusetts, USA), machines that are used routinely by *Drosophila* neuroscientists to study circadian rhythms and sleep. In particular, one of the most successful features of DAMs that we aimed to imitate is the ability to run dozens of experiments simultaneously, gathering data in real-time from thousands of flies at once, using a device that follows a “plug-and-play” approach. Here we describe the philosophy and technical vision underlying ethoscopes. We provide some examples of raw and processed data that users will be able to acquire, and offer some proof-of-principle examples of the feedback-loop stimulus.

## Results

### Principle, scope and availability

An ethoscope is a self-contained machine able to either record or detect in real-time the activity of fruit flies (and potentially other animals) using computerised video-tracking. It relies on an independent small single-board computer (Raspberry Pi[20] - rPi) and a high-definition camera (rPi camera[20]) to capture and process infrared-illuminated video up to a resolution of 1920x1080 pixels, at 30 frames per second (Fig 1A). Ethoscopes are assembled in a 3D-printed chassis and, with cables, they have an approximate footprint of 10x13x19 cm (Fig 1B and **S1 Interactive figure**). Although we recommend a 3D-printed assembly for research-grade use, we also provide detailed instruction to build a fully functional ethoscope out of LEGO bricks (Fig 1C, - LEGOscope **S2 Supporting information**) or out of folded cardboard (Fig 1D, - PAPERscope **S3 Supporting information**). These latter two options are particularly well suited for the purpose of education and outreach. In all cases, assembly of ethoscopes requires little technical skill. The technical drawings required to 3D-print and assemble an ethoscope, along with its software (Python code on a Linux instance) are released under the open source GPLv3 license and are freely available on the ethoscope website (https://lab.gilest.ro/ethoscope). A current version of the user manual, including building instruction, is also provided here as **S4 Supporting information**, while current snapshots of STL and image files are available on Zenodo[21]. The combination of consumer-grade electronics, 3D printing and free open source software results in a total cost of about €100 for each machine. Software is provided as source on a GIT repository and as self contained images that can be written either on SD cards to fit inside each Raspberry Pi, or on a CD to work as the controlling unit (“the node” in Fig 2A). Limited cost, combined with each ethoscope relying on its own computing power, allows for easy scaling of the entire platform.

**Figure 1.**
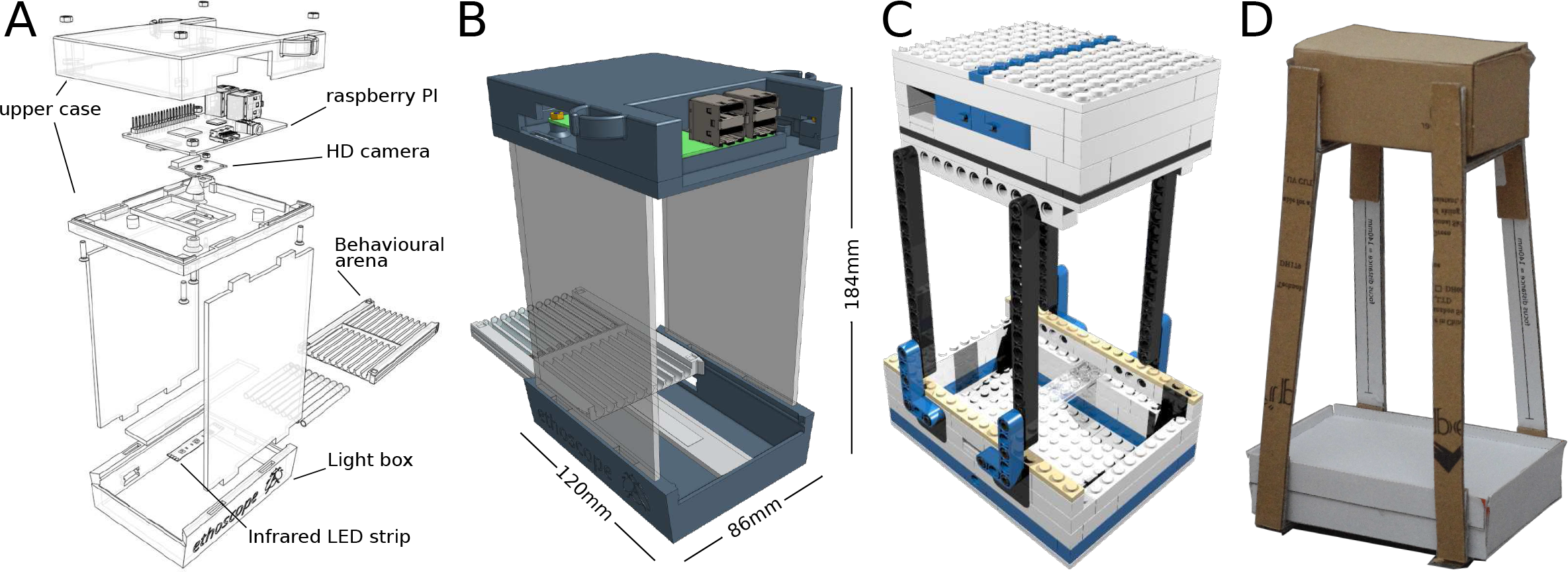
The ethoscope. (**A**) Exploded drawing of an archetypical ethoscope. The machine is composed of two main parts: an upper case housing the Raspberry Pi (rPi) and its camera, and a lower case providing diffused infrared light illumination and support for the experimental arena. The two cases are separated by spacers maintaining a fix focal distance (140 mm for rPi camera 1.0). (**B**) A rendered drawing of the assembled model, showing the actual size without cables. The presence of USB and connection cables will slightly increase the total size (cables not shown for simplicity). The arena slides in place through guides and locks into position. A webGL interactive 3D model is available as **S1 Interactive figure**. (**C**) The LEGOscope, a version of the ethoscope built using LEGO bricks. A detailed instruction manual is provided in **S2 Supporting information**. (**D**) The PAPERscope, a paper and cardboard version of the ethoscope, assembled using 220gsm paper and 1mm grayboard. Blueprints are provided in **S3 Supporting information**. In all cases, ethoscopes must be powered with a 5VDC input using a common USB micro cable either connected to the main or to a portable powerpack.

**Figure 2.**
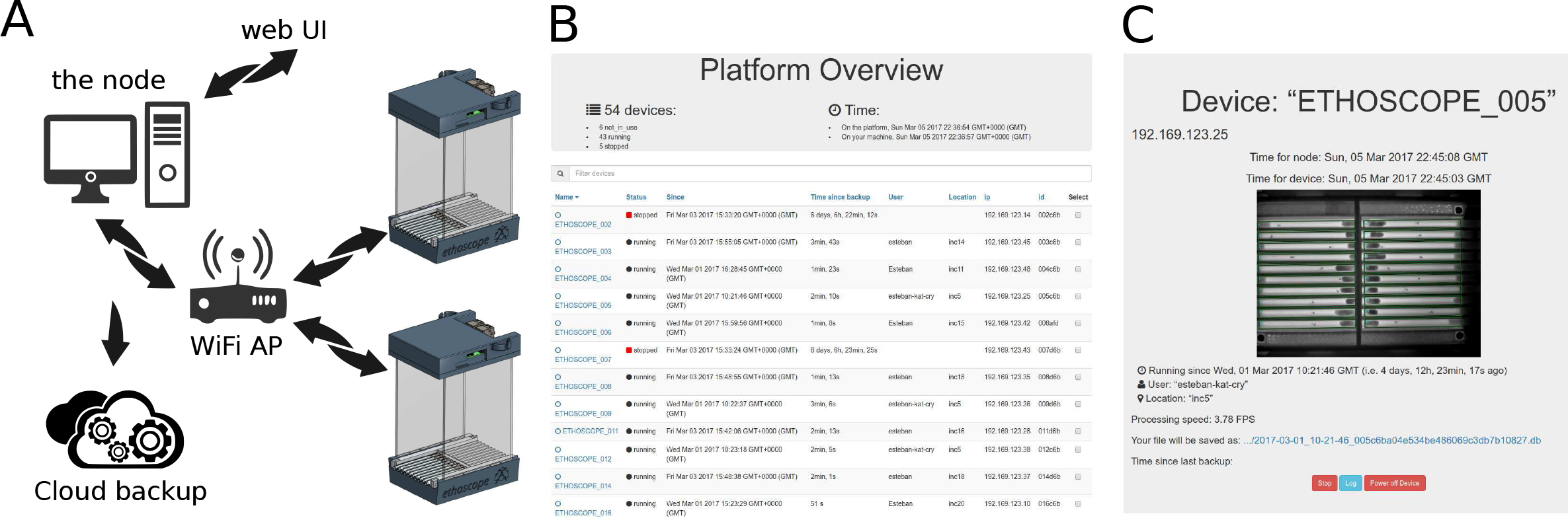
The ethoscope platform. (**A**) A diagram of the typical setup. Ethoscopes, powered through a USB adapter, are connected in an intranet mesh through an Access Point (AP) or a Wi-Fi router. A server computer in the network acts as “the node”, downloading data from ethoscopes and serving a web-based user interface (UI). Ethoscopes can be controlled through a web graphical user interface (web-UI), either locally or remotely. (**B**) Screenshot of the homepage of the web-UI, showing a list of running machines and some associated experimental metadata (e.g. username and location). (**C**) Screenshot of an ethoscope control page on the web-UI, providing metadata about the experiment and a real-time updated snapshot from the ethoscope point of view.

### Usage scenario

In a typical usage scenario, several ethoscopes are placed in a climate-controlled chamber. Each ethoscope is powered through a USB cable and communicates via Wi-Fi to a local network, uploading data to a desktop computer acting as the data collecting station (“the node” in Fig 2A). Through the same network, ethoscopes can be remotely commanded using a graphical web interface (Fig 2B,C and **S1 Video**). If the node is connected to the Internet, the entire platform will receive automatic software updates from the upstream GIT repository. Since each ethoscope operates independently, there is no theoretical limit to the number of machines that can be used concurrently. In fact, the ability to run dozens of ethoscopes simultaneously is one of the crowning features of the system. However, Raspberry Pis are quad-core microcomputers and they do generate considerable heat under heavy computing load. For this reason, the use of a climate-controlled chamber is a strict requirement and remains the greatest limitation of the platform at present. In our laboratory, we run up to 70 ethoscopes at once - analysing 1400 flies - spread across 20 commercial wine-coolers modified to be used as temperature controlled chambers (details of the modifications are available upon request). Besides being a good solution for multi-users environments, the use of many small climate chambers, rather than a few with greater capacity, also allows for more flexibility in designing and running experiments, for instance by running different cohorts at different temperatures for thermogenetic manipulation; or by running different time zones in the same room. Importantly, provided animals have access to fresh food, the platform is able to run experiments for weeks. Ethoscopes connected to the network will periodically transfer collected data to the node acting as a local storage server, meaning the duration of the experiment is not limited by the storage capabilities of the rPi.

### Behavioural arenas

The experimental flies are loaded into a behavioural arena that slides and locks inside the lower part of the ethoscope chassis (Fig 1A). Like the rest of the machine, arenas are 3D-printed and their design depends on the nature of the experiment: some examples of arenas inspired by commonly used behavioural paradigms are provided in Fig 3 and span arenas adopted for long-term experiments that may last for weeks, such as sleep or longevity analysis (Fig 3A-C,F), or short term assays such as decision making (Fig 3D) and courtship (Fig 3E,G,H). Arenas feature three fixed recognition marks on the corners (red circles on Fig 3A), that are used by ethoscopes to automatically align and register the regions of interest for tracking. When starting an experiment, the experimenter can decide whether the activity of the animals should be tracked in real-time or whether the ethoscope should record a video to be analysed offline, with the ethoscope software or with other software, such as the C-trax/JAABA suite[14,15], CADABRA[17] or idTracker[16]. In real-time tracking mode, ethoscopes will detect and record the position and angle of each animal with a variable frame rate that fluctuates between one and four FPS, depending on the computing load (e.g. number of flies to be tracked and number of regions of interest; see **S5 Supporting information** for technical details of real-time tracking and its performance).

**Figure 3.**
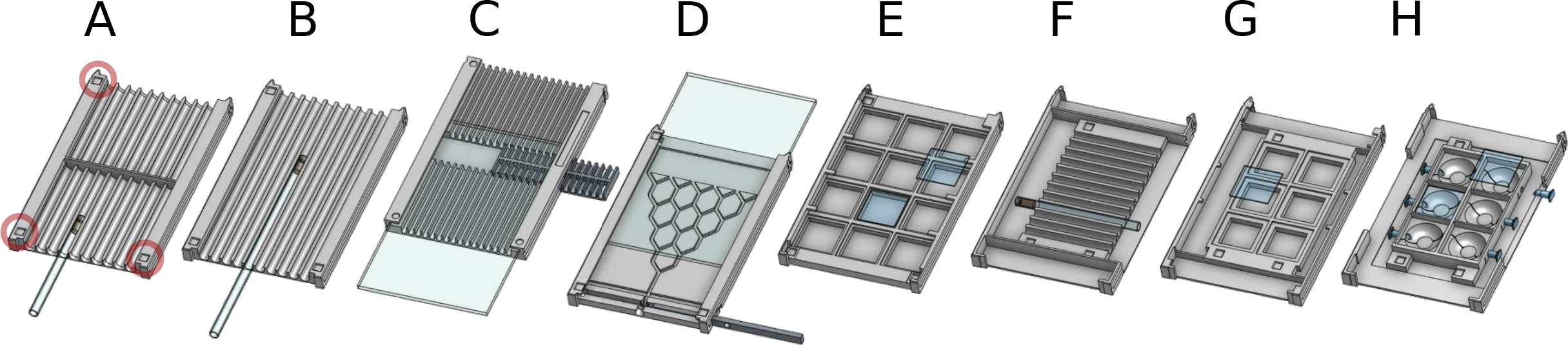
Versatility of use with custom behavioural arenas. (**A-H**) Examples of 8 different behavioural arenas whose files for 3D printing are available on the ethoscope website. (**A**) Sleep arena. Most commonly used arena for sleep studies, lodging 20 individual tubes. (**B**) Long tubes arena. Houses 13 cm tubes and can be used for odour delivery studies or, more generally, for behaviours requiring more space. (**C**) Food bullet arena. Animals are placed directly on the arena and food can be replaced by pushing in a new bullet[11]. Does not require glass tubes and can be used for quick administration of chemicals in the food. (**D**) Decision making arena. Can be used to study simple decision making behaviours - adapted from Hirsch[3]. (**E**) Square wells arena. Can be used for courtship assay or to record activity in a bi-dimensional environment. (**F,G**) Conceptually analogous to **A.** and **I.**, but designed to work in high-resolution (full-HD) settings. (**H**) Round wells arena, modelled following specifications from Simon and Dickinson[43]. Note that all arenas are marked with three visible reference points (indicated by a red circle in **A.**) that are used by the ethoscope to automatically define regions of interest for tracking, providing a degree of physical flexibility.

### Real-time tracking and behavioural profiling

The ethoscope software is modular in design, meaning many components can be replaced or adapted as needed. The tracking module is one that the end users may want to adapt to their needs ultimately. Currently, we provide two tracking options: an adaptive background subtraction model (default option - **S5 Supporting information**) and an experimental tracking module based on haar-cascades[22], that is suitable for tracking multiple animals in the same region of interest without maintaining their identities. To validate the accuracy of the default tracking mode, we asked three experienced fly researchers to manually annotate the position of the flies in 1413 still frames extracted from 2736 hours of recorded videos. We then compared the manually annotated positions to the coordinates of the fly centroids as detected by the ethoscope tracking software, and found a strong degree of overlap, with a median discrepancy of 300 μm, corresponding to a tenth of a fly body length. In no cases (0/1413 frames), did the error exceed one body length (2.5 mm). To enrich the capabilities of ethoscopes, we also implemented a real-time behavioural annotator. We created a ground-truth of 1297 videos, each lasting 10 seconds and each manually annotated by at least three experienced fly researchers (Fig 4A, annotation labels were: “walking”, “micro-movement” or “immobile”). Random forest variable importance[23] was used to screen for predictors of movement in a supervised manner and the two highest-ranking features - maximal velocity and cumulative walked distance - were selected for further analysis. Conveniently, maximal velocity alone appeared to serve as a faithful predictor of behaviour (Fig 4B) allowing for real-time dissection of basic behaviour. Therefore, not only can ethoscopes reliably annotate the position of flies, but also detect when an animal is immobile, performing a micro-movement (such as grooming, eating, and egg laying), or walking, with an accuracy of 94.3% for micro-movement detection and 99.0% for walking detection. As proof of principle, we show a low resolution (5 days with a definition of 30 minutes - Fig 4C) and a medium-resolution (3 hours with a definition of 10 seconds - Fig 4D) activity plot for 10 individual animals (5 young males and 5 young females, between 4 and 9 days old).

**Figure 4.**
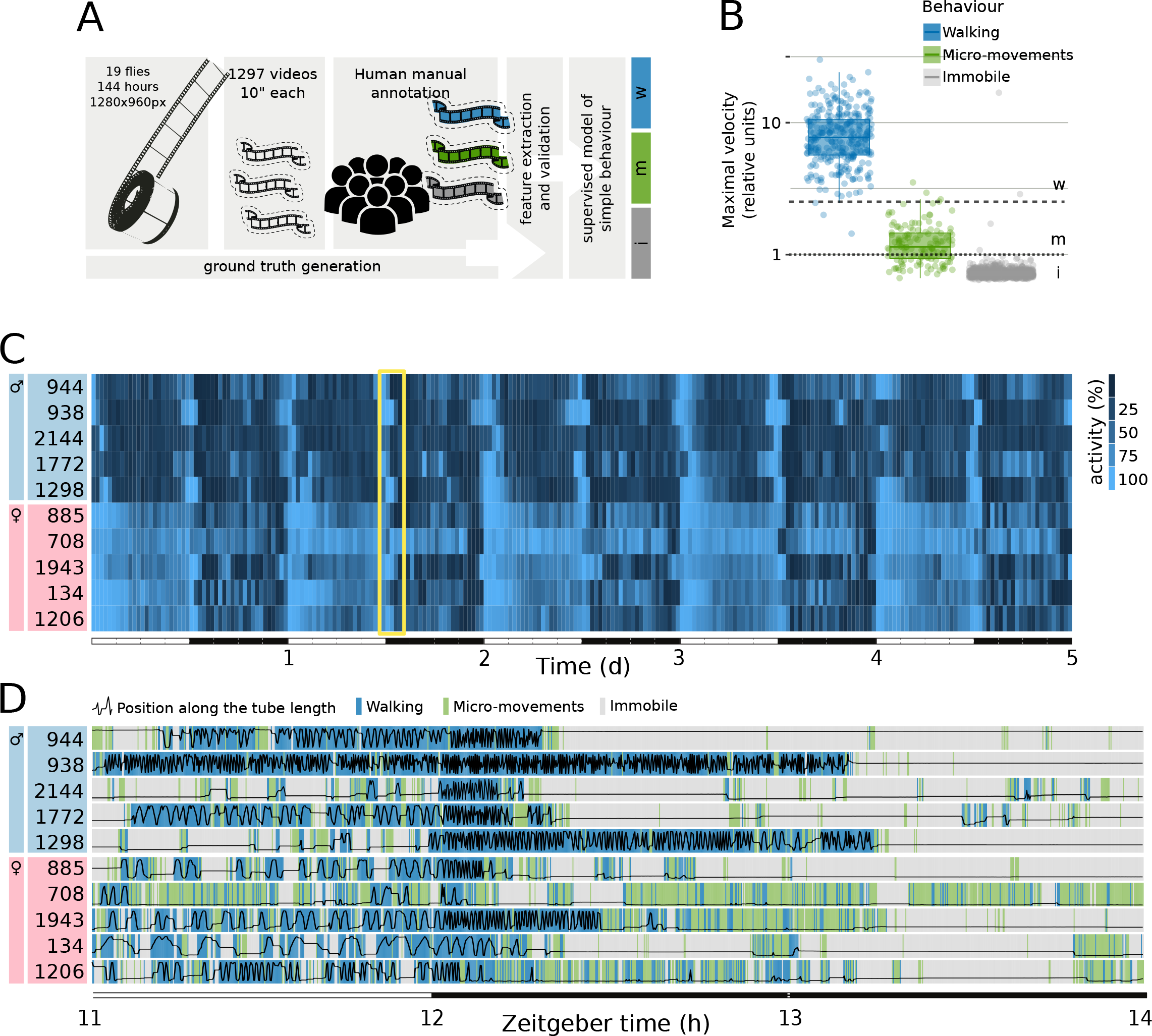
Tracking and validation of behavioural classification. (**A**) To build a statistical model of activity, we used ethoscopes to record offline 2736 hours of video (144 hours x 19 flies) at resolution of 1280x960 pixels and frame rate of 25 FPS. Video fragments of the duration of 10 seconds were sampled every hour for all 19 animals and scored by at least three experienced fly researchers in a randomised order. Consensual annotations - where majority of scorers agreed - were kept, resulting in a ground truth of 1297 video fragments (116 ambiguous annotations were excluded using this latter criteria). Scorers manually annotated both the position of the animal in the tube and the perceived behavioural state (i.e. immobile, micro-moving or walking). Ethoscope video-tracking was run independently on the whole video down-sampled between 1 and 5 FPS, all realistic frame rates for real-time analysis. (**B**) Distribution of corrected maximal velocity (relative unit, see **S5 Supporting information**) for each behaviour, showing the thresholds used to detect movement (1-dotted line) and walking (2,5 - dashed line). (**C**) Five days recording of activity of ten representative flies: 5 males (cyan boxes) and 5 females (rose boxes). Flies were kept in a regime of constant climate, in a 12h:12h light-dark cycle (as indicated by the lower bar alternating white and black). The yellow frame highlights the three hour window shown in **D**. (**D**) Detailed activity for the same individuals shown in **C**., during a three hour window spanning a light to dark transition. The black line shows the position of the animals from the food end to other extremity of the tube (bottom to top). The background colours highlight the behavioural features as detected in real-time by the ethoscope, with a definition of 10 seconds per pixel (same legend as **B**).

### Real-time feedback-loops on single animals

The ability to operate in real-time offers a crucial feature: animal-specific feedback-loop stimuli delivered upon a predefined behavioural trigger. Interfering with the behaviour of an animal through external stimuli is an important tool for neuroscientists. In principle, feedback-loops can be used for multiple purposes: to reinforce learning, to sleep deprive animals, to stimulate or silence circuits using optogenetics, to study operant conditioning, *et cetera*. Systems operating feedback-loop stimuli on fruit flies have been proposed previously and have already proved to be instrumental, but are not easily compatible with a high-throughput approach and are focussed on very specific usage[24,25]. We therefore designed ethoscopes so that they could be extended with modules that seamlessly connect with the machine and react in real-time to trigger an action whenever a condition is satisfied. Figure 5 shows three examples of such modules: an air/gas/odour (AGO) delivery module (Fig 5A,B), a rotational module (Fig 5D,E), and an “optomotor” module combining optogenetic stimulation and motor disturbance (Fig 5G,H). All modules plug into the bottom part of the machine and are configured through the main graphical web-interface, where the experimenter can set the trigger conditions that will activate the stimulus and schedule a time window for their function (**S1 Video**). A trigger can be a combinatorial ensemble of position, time, and behaviour (e.g. *“micro-movement for at least 20 seconds within 5mm from the food” or “immobile for at least 5 minutes anywhere*”). As proof of principle, we provide representative evidence of how single flies react to three different stimuli: a 5 second delivery of CO_2_, triggered by crossing of the midline tube (Fig 5C); a 2 second fast rotation of the tube (60° / 0.12 seconds), triggered by 20 seconds of immobility (Fig 5F); a 5 second opto-stimulation on “moon-walker”[26] receptive flies, manually or automatically triggered (**S2 Video**). We also provide a case test for using the rotation module as a sleep-deprivation device (Fig 5I-P). To this date, scientists studying sleep in flies have the option of performing mechanical sleep deprivation by placing animals on an orbital shaker[27], a rotating device[28], or a vibrating platform[10]. In all cases, the resulting mechanical stimulation of the animals is independent of their actual activity, so that the stimulus is delivered unspecifically to all individuals at the same time: to some while asleep, and to others while awake. Using this module, we can rotate single tubes - hence, single animals - only when a fly is actually immobile (e.g. after 20 consecutive seconds of immobility, Fig 5I-L) or, in the yoked control, only when a fly is actually walking but not eating or grooming (e.g. after midline crossing, Fig 5M-P). A conceptually identical paradigm was originally introduced in the 1980s[29] and it is still considered one of the best controlled paradigms for chronic sleep deprivation of rodents. As shown in Figure 5, all flies are subjected to an analogous number of tube rotations (548 ± 342 for experimental sleep deprivation; 383 ± 173 for yoked control; mean ± SD-temporal pattern shown in Fig 5K,O), but only the experimental sleep deprivation leads to a sleep rebound after the treatment (Fig 5L,P), thus confirming that sleep rebound is indeed a specific counter-effect of sleep deprivation[30]. For sleep scientists, the possibility to precisely deprive flies of sleep may be a crucial tool to differentiate the effects of mere sleep deprivation from the effects of stress, two confounded phenomena[28]. On the ethoscope website, we provide detailed instruction on how to build all three modules in conjunction with a description of the API needed to interface any new custom module to the ethoscope platform.

**Figure 5.**
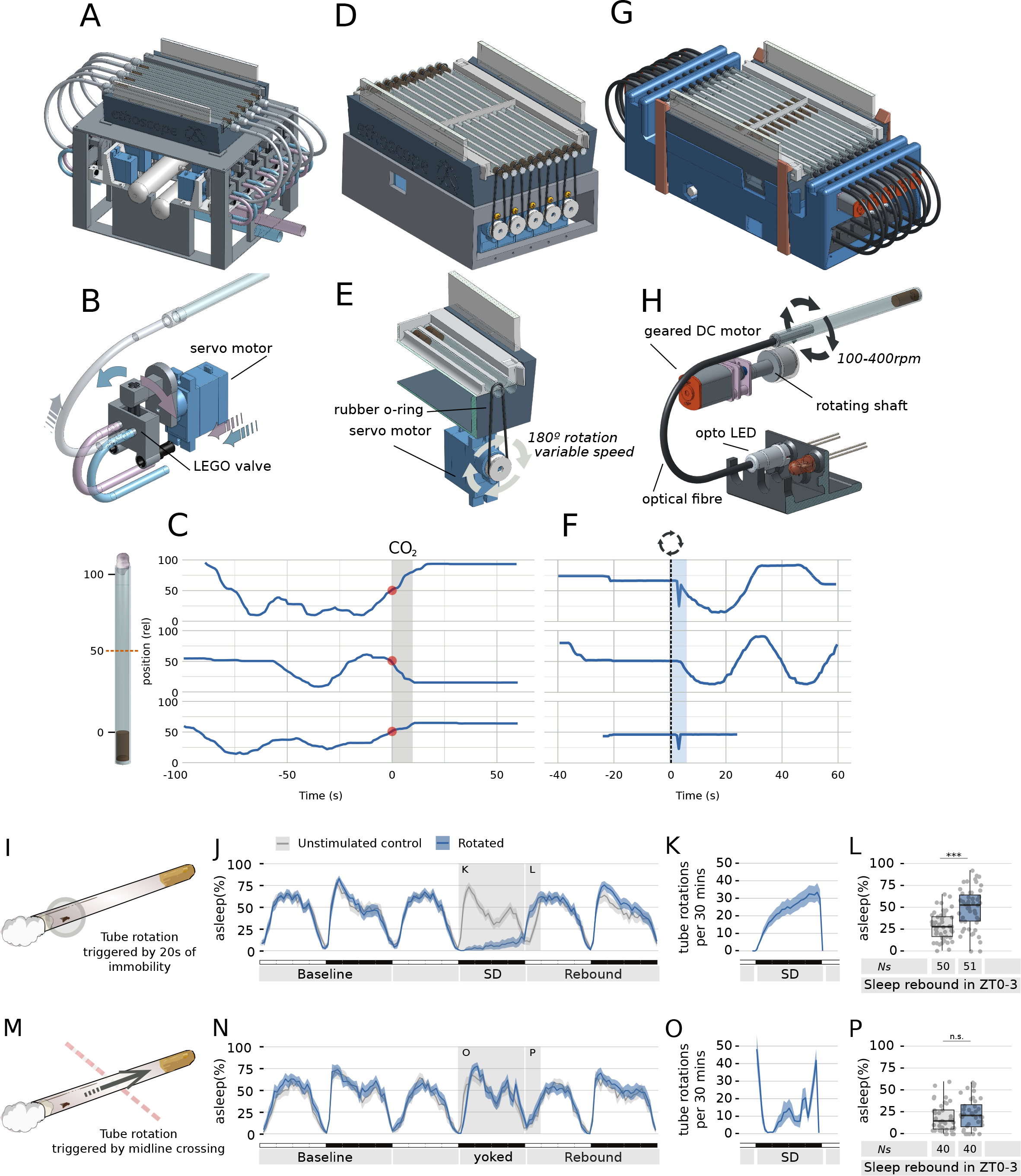
Versatility of use with behavioural feedback-loop modules. (**A**) Diagram and (**B**) detail of the air/gas/odour (AGO)-delivery module. Two independent flows (blue and purple in the drawing) are fed into the module using external sources. The module features 10 LEGO valves, each independently controlled through a servo motor. The motor switches the air source on the valve, selecting which source will be relayed to the tube containing the fly. Available positions are: blue source, purple source, and closed. (**C**) Representative response of three flies subjected to CO_2_ administration using the AGO module. CO_2_ release lasts 5 seconds (grey bar) and it is triggered by midline crossing (red dot). The blue line indicates the fly position in the tube over the 150 second period. (**D**) Model and (**E**) detail of the rotational module. The module employs a servo motor to turn the tube hosting the fly. Direction, speed, duration and angle of the rotation can be modulated to change the quality of the stimulus. (**F**) Representative response of three flies upon stimulation using the rotational module shown in (**D,E**). Rotation of the tube is triggered by 20 consecutive seconds of immobility (dashed line) and is followed by 5 seconds of masking during which tracking is suspended to avoid motion artefacts (cyan area). The bottom panel shows traces of a dead fly. (**G**) Model of the optomotor module, able to simultaneously stimulate single flies with rotational motion and light. (**H**) Detailed view of the optomotor principle. Light is directed into the tube using optical fibre. **S2 Video** shows the optomotor module in action. (**I-P**) The servo module employed for a sleep deprivation experiment. Flies shown in grey are unstimulated mock controls, never experiencing tube rotations. Flies shown in light blue experience rotation either after 20 seconds of inactivity (**I-L**) or after midline crossing (**M-P**). (**J,N**) Sleep profile of flies along three days in conditions of 12h:12h light dark cycle. Gray shadings indicated the stimulation period and the following sleep rebound period. (**K,O**) number of tube rotations delivered during the 12h stimulation period. (**L,P**) Quantification of sleep rebound during the first three hours of the day following the stimulation (ZT: zeitgeber time).

## Discussion

### Strengths and limitations

Ethoscopes emerge from the maker culture and combine three important innovations of the last decades-3D printing, small single-board computers and machine learning - into a novel tool for behavioural researchers. They were designed to be easy to build, inexpensive, and compatible with high-throughput research. Accessibility and high-throughput design are certainly two important features of the platform but we anticipate that the combination of those two with the ability to create custom feedback-loop experiments, will make ethoscopes particularly useful for the community. Creating feedback-loop based experiments is something that *Drosophila* neuroscientists have been doing for decades, with great ingenuity and success[24,31–33]. However, these generally require *ad-hoc* equipment and provide limited procedural throughput. Ethoscopes build upon this tradition but offer a modular platform that may simplify this procedure and favour wide adoption.

The philosophy of distributed microcomputing is one of the strongest features of the ethoscope platform - in terms of affordability and scalability - but at the same time it constitutes its current greatest weakness: relatively limited computational power. In their current form, ethoscopes rely on Raspberry Pis and work best when sporting their third, most powerful, hardware version (rPi 3). In principle, however, any microcomputer platform able to connect to a camera would work and it is possible that future versions may take advantage of commercial development to improve computational power and, ultimately, performance. As of now, real-time tracking is limited to a temporal resolution of 1-4 Hz. Whenever greater temporal resolution is needed, the offline tracking mode transforms ethoscopes into “remotely controlled video cameras” and allows users to acquire video files at up to 90 frames per second, to be analysed at a later stage with another software of choice. If greater spatial resolution is needed, it is possible to expand the rPi cameras with lenses featuring an M12 mount. The possibility of coupling rPi cameras to lenses has been demonstrated recently by the FlyPi project, a tool very similar in philosophy but different in scope[34]. The fruit fly community has produced excellent software for automatic recognition of complex behaviours[14,16,35] with the demonstrated potential of revolutionising the field[36]. Ethoscopes can contribute and assist to this end, by facilitating scalability.

### Future developments and improvements

We anticipate that one of the most interesting development of the platform may be the growing variety of feedback loop modules. Here we offered three examples of feedback-loop modules that can be used to expand ethoscopes’ abilities and, ultimately, we expect and encourage users to build modules based on their own needs, increasing the available range of modules. For instance, scientists studying feeding behaviour may want to add a module able to simultaneously record access to food, using the expresso[13] or flyPAD[12] technologies; use of collimated high-power LEDs coupled to small optical filters could also be used to create a module for visualisation of immunofluorescence in real-time[34]. Another possible future improvement may derive from the announced introduction of machine-learning dedicated chips (TPUs), currently being developed by tech giants such as Google, Microsoft and NVidia. It is likely that future versions of microcomputers will possess some form of TPUs, and that may allow for much more powerful discrimination between behaviours in real time.

Another possible use of ethoscopes is the adaptation of the platform to detect behaviour of other animals. Clearly, adapting ethoscopes to work with other small insects similar to Drosophila should be an easy task; tracking behaviour of even smaller animals may be possible using lenses and, for larvae or worms, modified illumination techniques, such as FTIR[37–39].

## Methods

### Model design and 3D printing

All parts were designed using the web SaaS onshape (http://www.onshape.com). All components were printed using Ultimakers 2+ (Ultimaker, Geldermailsen, Netherlands), with 2.85 mm PLA filament (RS 832-0273). STL to gCode translation was achieved using Cura (https://github.com/Ultimaker/Cura).

### Electronics

Electronic components were obtained through RS components, UK and Farnell, UK. A complete up to date bill of materials is available on the ethoscope website and here in **S4 Supportive information**.

### Data analysis and statistics

All data analysis was performed in R[40], using the R package Rethomics (https://github.com/gilestrolab/rethomics) and statistical analysis (Fig. 5L,P) consisted of pairwise Wilcoxon rank sum test (i.e. Mann-Whitney U test). For the sleep plots (Fig. 5J,K,N,O), bootstrap re-sampling with 5000 replicates, was performed in order to generate 95% confidence interval[41] (shadowed ribbons around the mean in the figures). Ns indicate the total number of flies over all experiments. Statistics were performed on aggregated data. Outliers were never excluded. Flies that died during the course of the experiment were excluded from all analysis. Traces and plots were generated in R, using ggplot2[42]. For all the boxplots, the bottom and top of the box (hinges) show the first and third quartiles, respectively. The horizontal line inside the box is the second quartile (median). Tuckey’s rule (the default), was used to draw the “whiskers” (vertical lines): the whiskers extend to last extreme values within +- 1.5 IQR, from the hinges, where IQR is Q3-Q1.

## Acknowledgements

The research leading to these results has received funding from BBSRC through BB/M003930/1, BB/J014575/1, and “Imperial College London - BBSRC Impact Acceleration Account”. EJB was supported by EMBO ALTF 57-2014 and by the People Programme (Marie Curie Actions) of the European Union’s Eighth Framework Programme H2020 under REA grant agreement 705930. We thank the Imperial College London Advanced Hackspace (ICAH) for granting early access to their equipment. Special thanks to Stefanos Zafeiriou and Susan Parker for technical discussions, to Anne Petzold for setting up the optogenetics crosses, to Anya Battle Lindstrom for designing the rounded wells courtship arena. The UAS-CsChrimson:mCherry line was generated by the Jayaraman laboratory (HHMI) and the moonwalking VT50660-Gal4 line was a gift of Barry J. Dickson (HHMI).

## Authors’ contributions

QG, LGR and GFG designed the platform; QG and LGR wrote the software; QG and EJB performed the experiments; ARJ contributed the optomotor module; ASF contributed the AGO module; all authors contributed the manuscript.

## Competing interests

The authors declare no competing conflict of interest.

**S1 Interactive figure** | Interactive 3D rendering of the assembled ethoscope - requires webGL capable browser (e.g. Google Chrome)

**S2 Supporting information** | Instruction booklet for building a LEGOscope

**S3 Supporting information** | Instruction booklet for building a PAPERscope

**S4 Supporting information** | User manual and instruction manual for the ethoscope

**S5 Supporting information** | Technical description of the tracking algorithm

**S1 Video** | An overview of how the ethoscope platform works

**S2 Video** | The optogenetics component of the optomotor module in action. Moonwalking flies (VT50660-Gal4 :: UAS-CsChrimson) are illuminated for 5-7 seconds using a red LED (630nm) through an optical fibre. Illumination is either manually triggered (first part of the video), or triggered by the fly position.

